# Improving QSAR Modeling for Predictive Toxicology using Publicly Aggregated Semantic Graph Data and Graph Neural Networks

**DOI:** 10.1101/2021.08.08.455550

**Authors:** Joseph D. Romano, Yun Hao, Jason H. Moore

## Abstract

Quantitative Structure-Activity Relationship (QSAR) modeling is the most common computational technique for predicting chemical toxicity, but a lack of methodological innovations in QSAR have led to underwhelming performance. We show that contemporary QSAR modeling for predictive toxicology can be substantially improved by incorporating semantic graph data aggregated from open-access public databases, and analyzing those data in the context of graph neural networks (GNNs). Furthermore, we introspect the GNNs to demonstrate how they can lead to more interpretable applications of QSAR, and use ablation analysis to explore the contribution of different data elements to the final models’ performance.

## 1. Introduction

Evaluating the toxicity of chemicals is an essential component of pharmaceutical and environmental research. Traditionally, the task of establishing toxicity has been accomplished using *in vivo* models, where a model organism is exposed to a chemical of interest and observed for toxic effects, or by performing epidemiological studies on human populations. Both of these approaches are costly and time consuming,^1^ and given the hundreds of thousands of compounds of toxicological interest, innovative alternatives are needed to rapidly screen chemicals. In recent decades, predictive toxicology and large-scale chemical screening efforts have emerged to address this issue.^2,3^

Quantitative Structure-Activity Relationship (QSAR) modeling is arguably the most prevalent method for predicting *in silico* whether a chemical will cause a toxic response.^4^ Briefly, QSAR modeling involves collecting regularly structured quantitative descriptions of molecular structures (known as *fingerprints*), and then fitting a statistical model (e.g., logistic regression, random forest, etc.) to sets of chemicals where a toxic endpoint of interest is already known.^5,6^ Since each data point used to train a model is itself the outcome of a single experiment, QSAR is a meta-analysis approach that is complicated not only by the challenge of capturing relevant structural features of chemicals, but also by errors, biases, and ambiguities in the underlying experiments used to generate the training data. Consequently, QSAR is often criticized for its disappointing performance on many tasks.^7,8^ The computational toxicology community has long acknowledged the need for new methodological innovations to improve QSAR performance, but few have been effectively implemented.

In this study, we address these issues by augmenting the traditional QSAR approach with multimodal graph data aggregated from several public data sources, and analyzing those data in the context of a heterogeneous graph convolutional neural network (GCNN) model. We evaluate the model using 52 assays and their accompanying chemical screening data from the Tox21 data repository, and compare its performance to two rigorously defined traditional QSAR models consisting of random forest and gradient boosting classifiers. Our results show that the GNN strategy significantly outperforms traditional QSAR. We further refine our results by removing various components of the graph to explain the relative contributions of different data sources to the GNNs’ increased performance. Finally, we discuss how GNNs improve the interpretability of QSAR, and suggest future directions to continue this body of work.

## 2. Methods

### 2.1. Obtaining toxicology assay data

We used the Tox21 dataset^2^—which is a freely available resource produced collaboratively by the US National Institutes of Health, the US Food and Drug Administration, and the US Environmental Protection Agency—to obtain a set of candidate assays for classification and establish ‘ground truth’ relationships between specific chemicals and those assays. Each assay in the database includes experimental screening results describing the activity of the assay in response to specific chemicals of toxicological interest, including pharmaceutical drugs, small molecule metabolites, environmental toxicants, and others. We removed all chemical–assay measurements with inconclusive or ambiguous results, as well as assays with very few (e.g., < 100) active chemicals.

### 2.2. Aggregating publicly available multimodal graph data

The graph data used in this study come from a new data resource for computational toxicology, named ComptoxAI^a^. ComptoxAI includes a large graph database containing many entity and relationship types that pertain to translational mechanisms of toxicity, all of which are sourced from third-party public databases (including PubChem, Drugbank, the US EPA’s Computational Toxicology Dashboard, NCBI Gene, and many others). We extracted the subgraph from ComptoxAI’s graph database defined as all nodes representing chemicals, genes, and toxicological assays, as well as the complete set of edges linking nodes of those types.

The 3 entity types that comprise the nodes of the extracted subgraph are *chemicals*, *assays*, and *genes*. We sourced the chemicals from the US EPA’s DSSTox database,^9^ and further filtered them so that each one is equivalent to a distinct compound in PubChem. We obtained genes from the NCBI Gene database,^10^ and assays from the Tox21 screening repository as described above. To serve as node features for chemicals, we computed MACCS chemical descriptor fingerprints^11^ for all chemicals in the graph, using their SMILES strings. Each fingerprint is comprised of a bit-string of length 166, where each bit indicates presence or absence of a specific chemical characteristic. These fingerprints are also reused as predictive features in the baseline (non-GNN) QSAR models, described below. All edges in the graph were sourced from either the Hetionet database^12^ or from assay–chemical annotations in Tox21. A metagraph describing the node and edge types in the subgraph is shown in **Figure 2**.

**Fig. 1.**
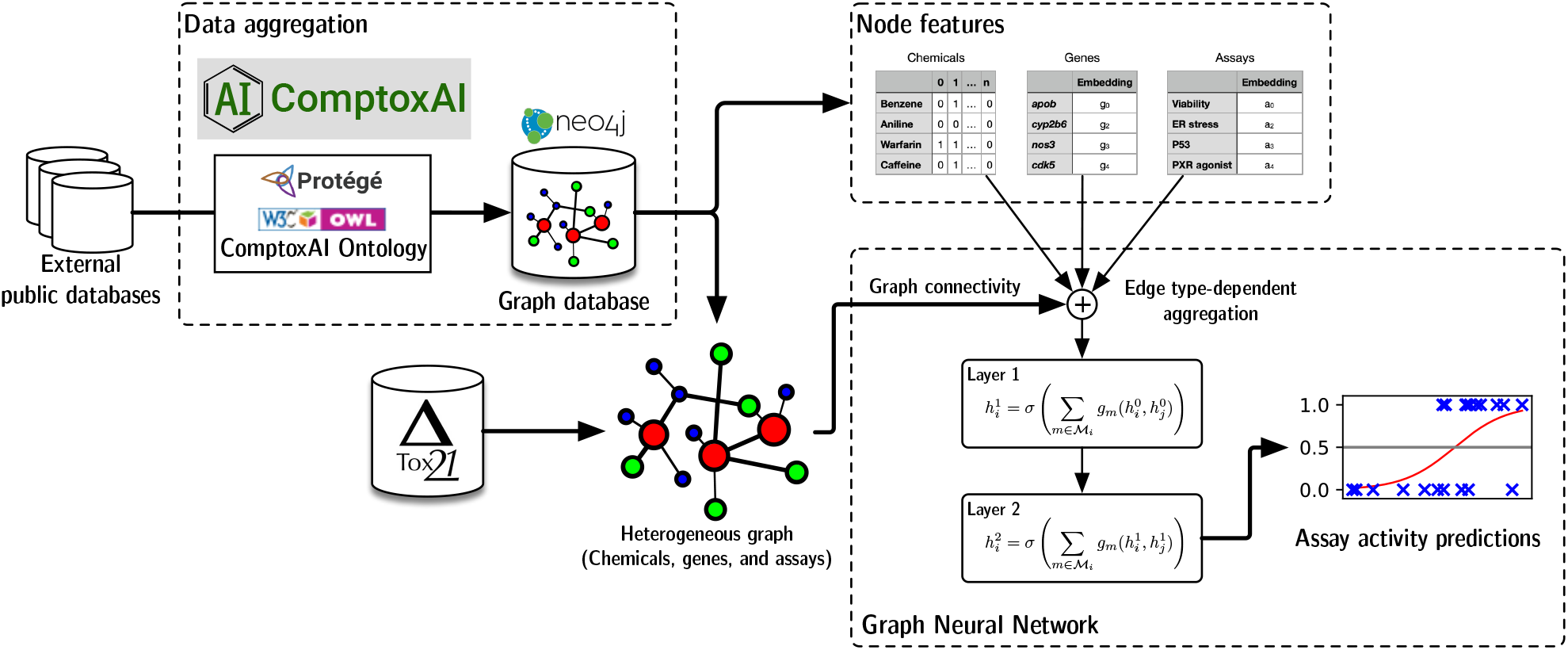
Overview of the graph machine learning approach used in this study. We build a toxicology-focused graph database (named ComptoxAI) using data aggregated from diverse public databases, and extract a subgraph for QSAR analysis containing chemicals, assays, and genes. We then train and evaluate a graph neural network that predicts whether or not a chemical activates specific toxicology-focused assays from the Tox21 database.

**Fig. 2.**
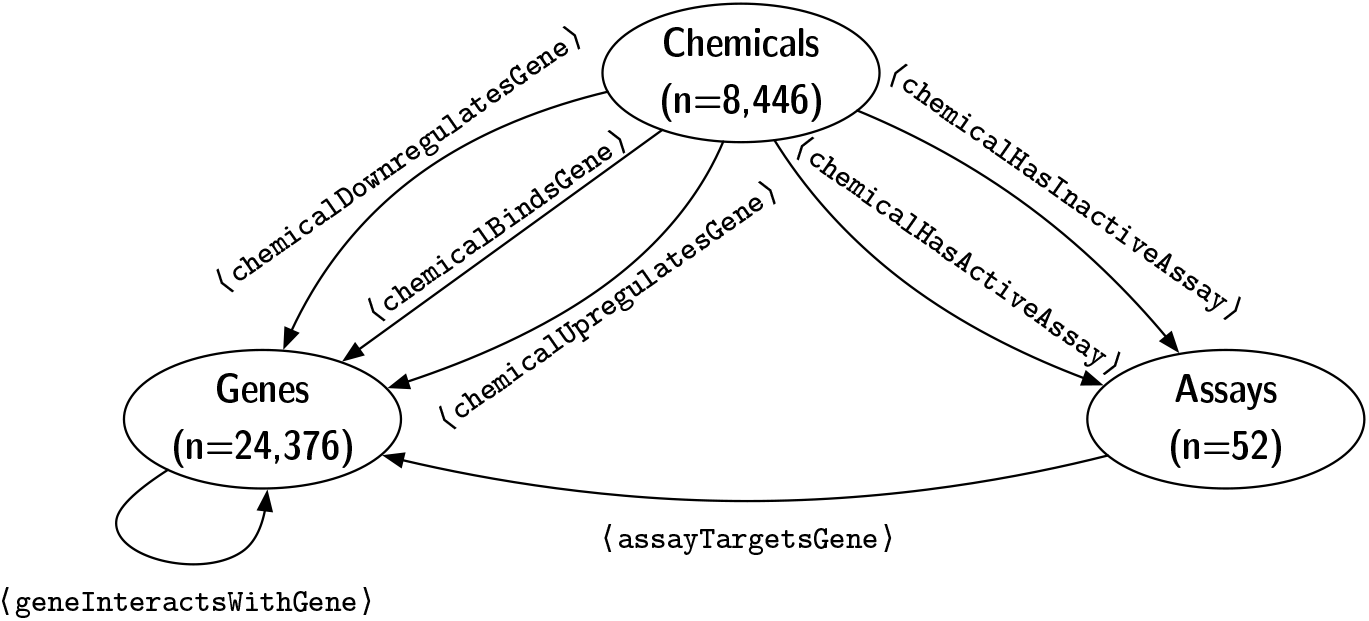
Metagraph describing the node types, node counts, and edge types in the heterogeneous graph. During implementation of the GNN, we also define corresponding inverse edges (e.g., assayTargetsGene ↔ geneTargetedByAssay) to facilitate the message-passing paradigm of the GNN.

### 2.3. Heterogeneous graph neural network

We constructed a heterogeneous graph convolutional neural network (GCNN) architecture^13^ for the graph ML experiments. Since our graph contains multiple entity types (chemicals, genes, and assays)—each with possibly different sets of node features, and linked by multiple semantically distinct edge types—the architecture extends the common GCNN model to learn separate message passing functions for each edge type. Briefly, each layer of the network aggregates signals from adjacent nodes in the graph, such that a greater number of layers results in signals being aggregated from an increasingly wider radius around each node. The output of the network can be thought of as encoded representations of nodes that incorporate information from the other nodes in their local neighborhood. The GCNN can also be thought of as a generalization of convolutional neural networks (CNNs) used in computer vision— instead of the convolutional operator aggregating signals from nearby pixels in an image, it aggregates features from adjacent nodes in the graph.^14^

In a heterogeneous graph, different node types represent different types of entities, each represented within a semantically distinct feature space.^15^ Therefore, the process of aggregating information from adjacent nodes must take those nodes’ types into account. Additionally, different edge types (e.g., ⟨`chemicalUpregulatesGene`⟩ and ⟨`chemicalDownregulatesGene`⟩) convey their own semantically distinct meanings, which can substantially effect the flow of information through the network. To handle these two challenges, we learn separate aggregation functions for each edge type in the graph, following the example proposed by Schlichtkrull *et al* in R-GCNs (Relational Graph Convolutional Networks).^16^ Within the R-GCN paradigm, the message passing process can be split into 3 sequential steps: (1.) collecting signals from adjacent nodes using an appropriate edge type-specific message function *ϕ*, (2.) combining each of those incoming signals (across all edge types) via a reduce function *ρ*, and (3.) finally updating the target node *v* by applying an update function *ψ*. Training the network is roughly equivalent to finding an appropriate parameterization of *ϕ* for each edge type.

A formal description of the GNN is given in **Appendix A**.

#### 2.3.1. Node classification

Given the GCNN architecture described above, we construct a heterogeneous graph where chemicals are labeled according to whether they do (1) or do not (0) activate an assay of interest. Although we remove the node representing the assay of interest^b^, all other Tox21 assays are included in the graph, and edges between chemicals and the other assays can therefore be used to improve the inferential capacity of the model beyond those of the baseline QSAR models, which only have access to chemical structure. Similarly, interactions involving genes further increase the information available to the model. We use the MACCS fingerprints as node features, while assay and gene nodes are initialized as 1-dimensional uniform random values that are optimized during model training, eventually serving as scalar ‘embeddings’ that are roughly proportional to those nodes’ importance in the trained network. The procedure we use for labeling the graph is shown in **Algorithm 1**.

**Algorithm 1.**
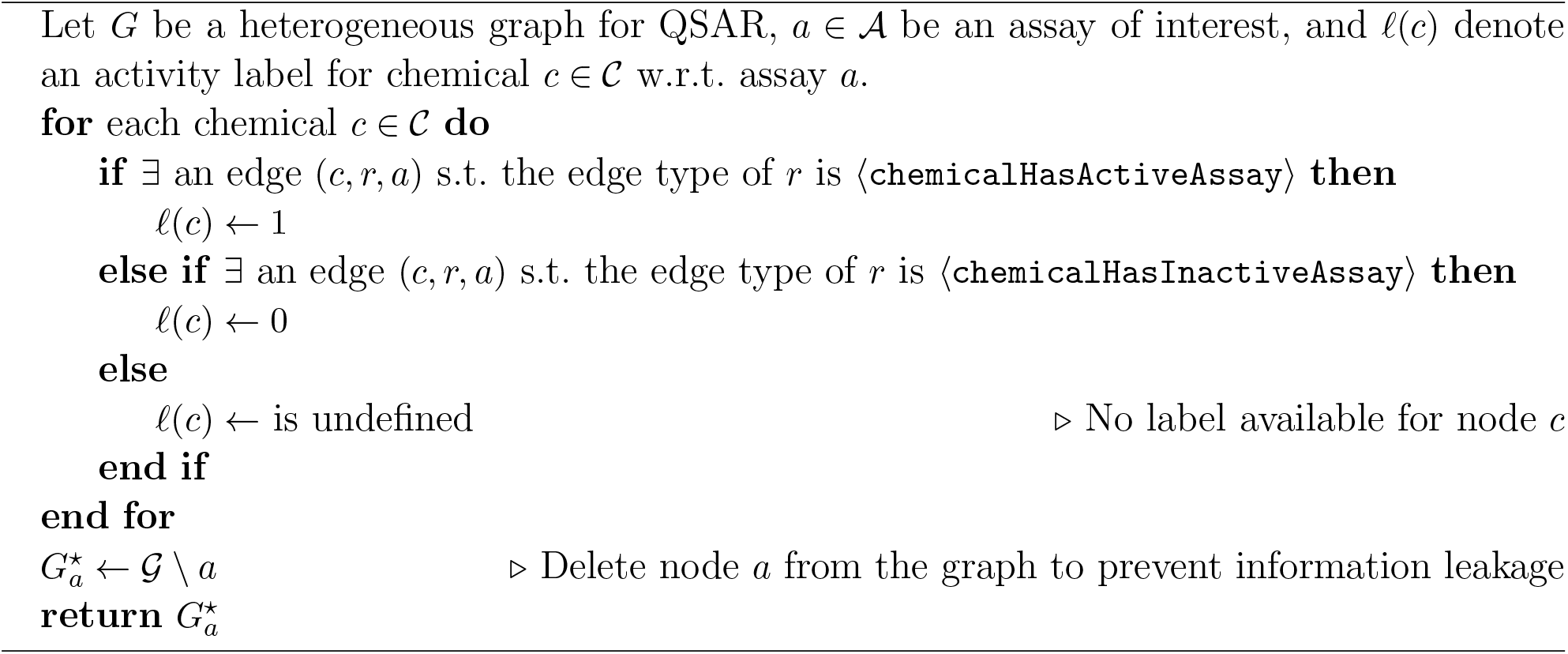
Labeled heterogeneous graph construction for toxicity assay QSAR model.

The resulting graph 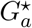 containing labeled chemicals is then used as input to the GCNN, which we train to predict the correct labels. We use an 80%/20% train/test split on the labeled chemicals, optimize the GCNN’s parameters using the Adam algorithm (a computationally efficient variant of stochastic gradient descent suitable for sparse gradients),^17^ and compute the error between predicted and true labels via cross entropy loss.

Additional details on the node classification approach are given in **Appendix B**.

### 2.4. Baseline QSAR classifiers

To assess the relative performance of the GNN classification model, we built 2 additional (non-NN) QSAR models that represent rigorously defined benchmarks consistent with current practice in predictive toxicology: A random forest classifier,^18^ and a gradient boosting classifier.^19^ Each model was trained on the aforementioned MACCS fingerprints of chemicals computed from SMILES strings, with an 80%/20% training/testing split. We tuned 6 hyper-parameters for each random forest model, and 5 for each gradient boosting model, as described in **Table S1**. These were tuned using grid search, where the optimal hyperparameter set is defined as the one that minimized binary cross entropy between predicted labels and true labels on the training data.

## 3. Results

### 3.1. GNN node classification performance vs. baseline QSAR models

Of the 68 total assays in the Tox21 database, we retained 52 for analysis in the QSAR experiments. The remaining 16 assays were not used due to either a low number of active chemicals or underrepresentation of screened chemicals in the ComptoxAI graph database. Additionally, we discarded compound labels for chemicals with inconclusive or ambiguous screening results.

As shown in **Figure 3**, the GNN model significantly outperforms both the random forest (Wilcoxon signed-rank test *p*-value 2.3 · 10^−4^) and gradient boosting (*p*-value 2.6 · 10^−3^) models in terms of area under the receiver operating characteristic curve (AUROC), with a mean AUROC of 0.883 (compared to 0.834 for random forest and 0.851 for gradient boosting). This is robust evidence that the GNN model tends to substantially outperform ‘traditional’ QSAR models. A notable characteristic of the GNN AUROCs is that their distribution has a higher variance than either the random forest or gradient boosting AUROCs. Anecdotally, this is likely due to diminished sensitivity of the GNN model when trained on assays with few positive examples—neural networks tend to struggle as data become more sparse, which seems to be the case here. We also compared F1-score distributions between the 3 model types; however, the differences between the 3 models are not statistically significant. The relatively low F1-scores in the 3 model types is a result of the class imbalance in the QSAR toxicity assays—all of the assays contain far more negative samples (assay is inactive) than positive samples (assay is active), which results in any false negatives having a magnified impact on F1. The same increased variance observed in GNN model AUROCs is shown in the GNN F1-scores.

**Fig. 3.**
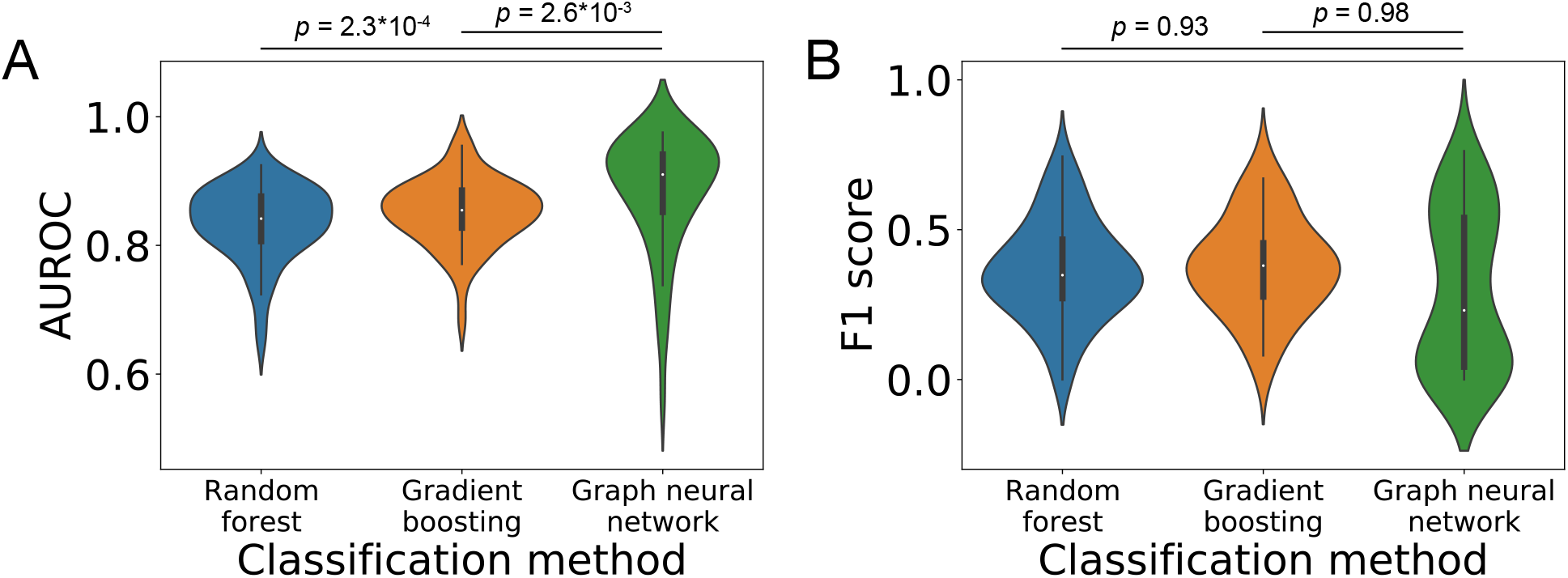
Overall performance metrics of the 3 QSAR model types on each of the Tox21 assays— a.) area under the receiver operating characteristic curve (AUROC) and b.) F1 score. The mean AUROC is significantly higher for the GNN model than for either of the two baseline QSAR approaches. The differences in F1 scores are not statistically significant. The GNN achieves poor F1 scores on assays with relatively few (e.g., < 100) “active” annotations in Tox21, which is consistent with known performance of neural networks on data with sparse labels. *p*-values correspond to Wilcoxon signed-rank tests on means, with a significance level of 0.05.

We performed further review of model performance on two selected assays of interest: PXR agonism (labeled `tox21-pxr-p1` in Tox21) and HepG2 cell viability (`tox21-rt-viability-hepg2-p1`). We selected these assays for two reasons: (1.) Both are semantically distinct from all other Tox21 assays (i.e., there are no other assays measuring pregnane X activity or cell viability), and therefore we would not expect an information leak from other highly correlated Tox21 assays present in the GNN, and (2.) both have a sufficient number of positive chemicals such that their ROC curves attain high resolution at all values of the decision rule across the 3 model types. **Figure 4** shows that the GNN outperforms the random forest and gradient boosting models at virtually all discrimination thresholds in both cases. The high performance of the GNN on HepG2 cell viability is especially noteworthy—cell viability is notoriously challenging to predict in chemical screening experiments. Many of the other 50 Tox21 assays showed similar patterns in performance. All ROC plots are available in **Supplemental Materials**.

**Fig. 4.**
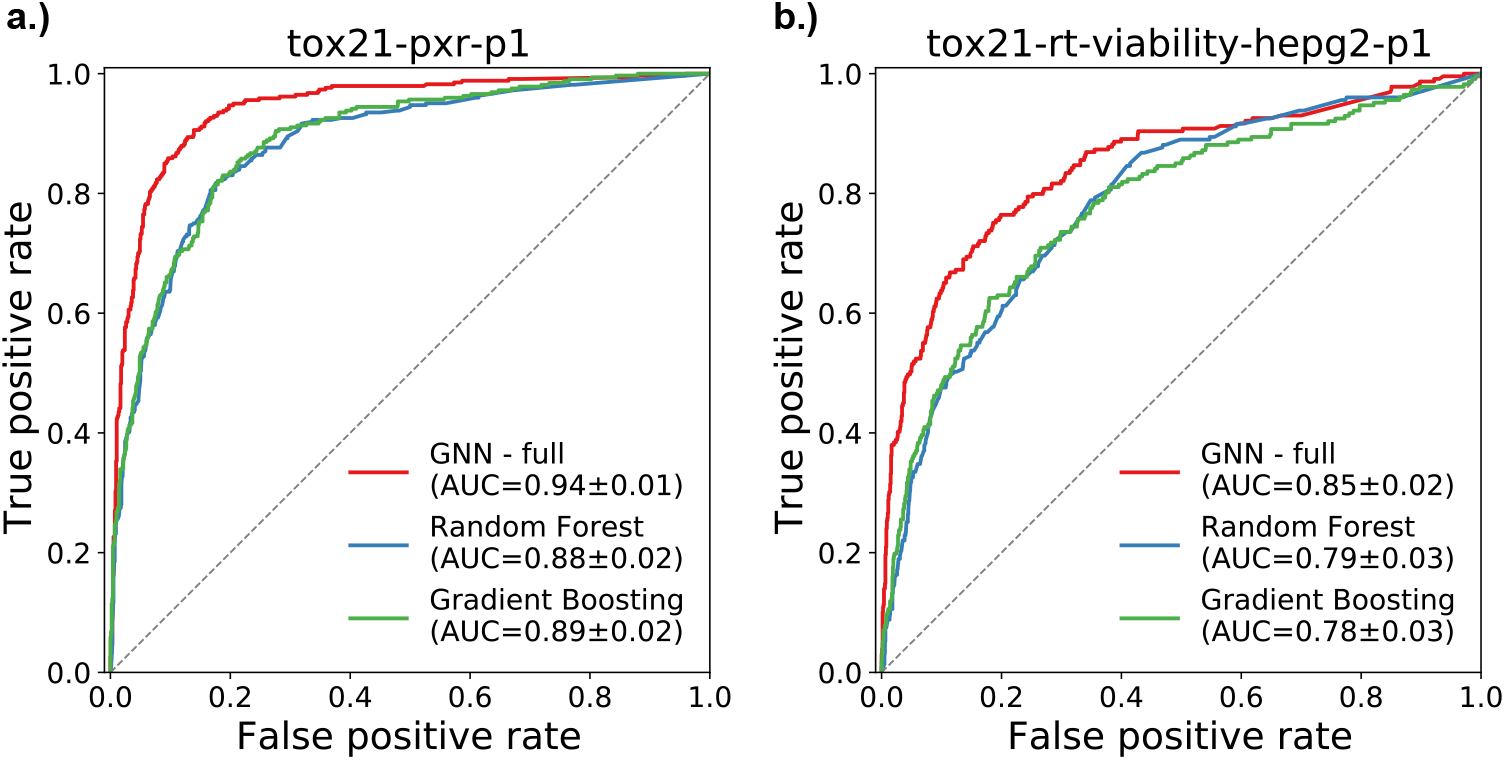
Receiver operating characteristic (ROC) curves for two selected Tox21 assays: a.) PXR agonism (`tox21-pxr-p1`) and b.) HepG2 cell viability (`tox21-rt-viability-hepg2-p1`). In both cases, the area under the curve (AUC) is significantly higher for the GNN model than either the Random Forest or Gradient Boosting models. AUC values are given with 95% confidence intervals.

### 3.2. Ablation analysis of graph components’ influence on the trained predictive model

To better understand how the GNN model outperforms the random forest and gradient boosting models, we performed an ablation analysis on the two previously mentioned assays— pregnane X agonism and HepG2 cell viability. For both of the assays, we re-trained the model after removing specific components from the GNN:

- All assay nodes.
- All gene nodes.
- MACCS fingerprints for chemical nodes (replacing them with dummy variables so the structure of the network would remain the same).

ROC plots for these experiments are shown in **Figure 5**. For both assays, the full GNN model performed best, although only modestly better (in terms of AUROC) than the versions without MACCS fingerprints or gene nodes. However, the performance of the GNN drops substantially—barely better than guessing labels at random (which would correspond to an AUROC of 0.50)—when assay nodes are removed from the graph. In other words, much of the inferential capacity of the GNN models are conferred by chemicals’ connections to assays other than the one for which activity is being predicted. Similarly, MACCS fingerprints are not—on their own—enough for the GNN to attain equal performance to the baseline QSAR models, which only use MACCS fingerprints as predictive features. Therefore, although the GNN achieves significantly better performance than the two baseline models, it is only able to do so with the added context of network relationships between chemicals, assays, and (to a lesser degree) genes.

**Fig. 5.**
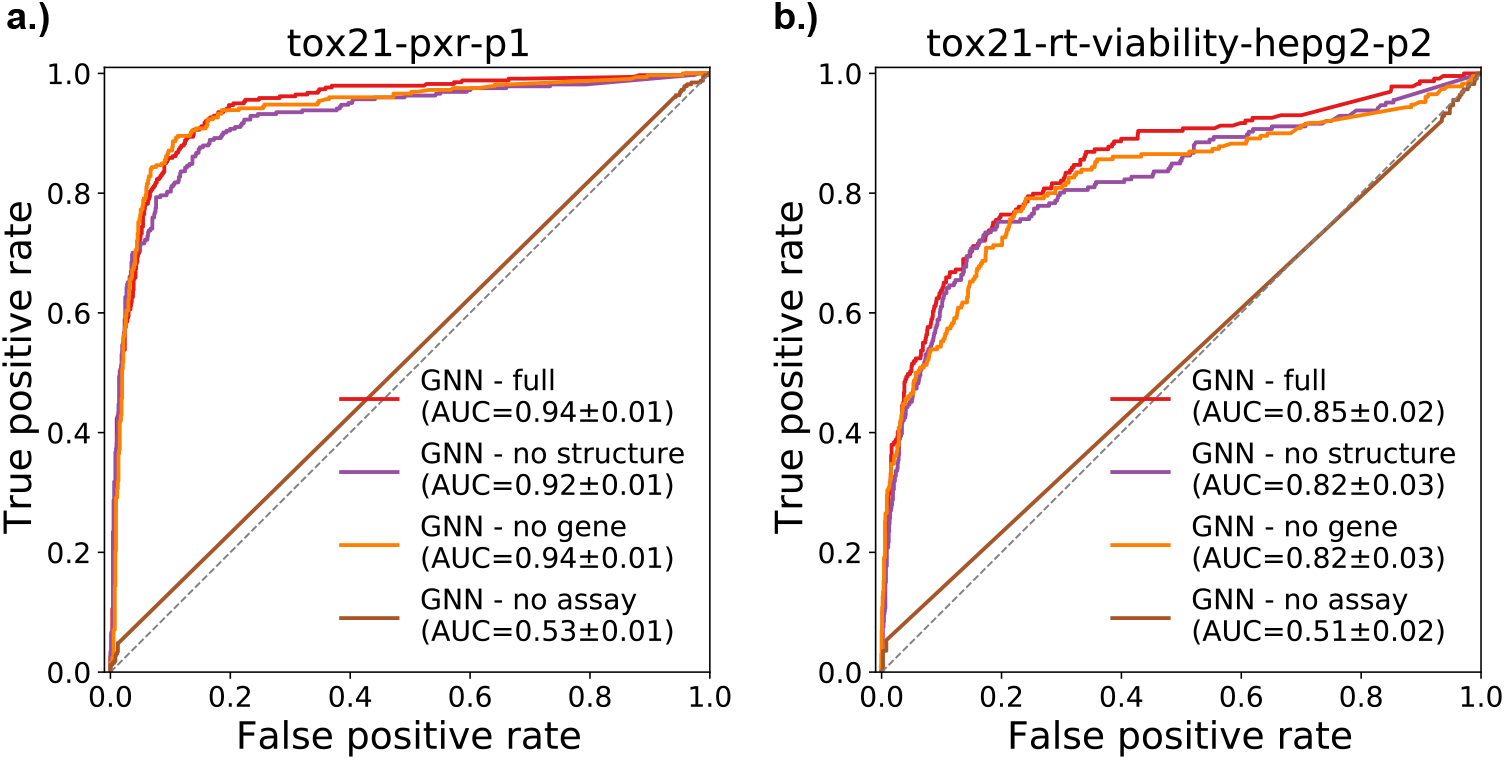
Receiver Operator Characteristic (ROC) curves for two selected Tox21 assays using different configurations of the GNN model. ‘GNN - full’ is the complete model as described in §2.3.1. ‘GNN - no structure’ omits the MACCS chemical descriptors and replaces them with node embeddings of the same dimensionality. ‘GNN - no gene’ removes gene nodes and their incident edges from the network. ‘GNN - no assay’ removes all assay nodes and incident edges, so predictions are made solely using chemicals, genes, the remaining edges, and the MACCS chemical descriptors as chemical node features. AUC values are given with 95% confidence intervals.

## 4. Discussion

### 4.1. GNNs versus traditional ML for QSAR modeling

The toxicology community largely agrees that QSAR underperforms on many tasks, and that new methodological advances are desperately needed. In this study, we demonstrate that GNNs significantly outperform the current gold-standard techniques in the field. Aside from the fact that neural networks can more easily adapt to nonlinear objectives than non-neural network models,^20^ this is likely a natural consequence of incorporating biomedical knowledge that goes beyond chemical structure characteristics. Gene interactions provide clues about how chemicals influence metabolic and signaling pathways *in vivo*, and non-target assays (i.e., other assays in the graph aside from the one currently being predicted) may correlate with activity of the target assay.

### 4.2. Interpretability of GNNs in QSAR

Chemical fingerprints—such as MACCS, which we use in this study—provide a valuable approach to representing chemicals that is suitable for machine learning. However, models based on fingerprints are challenging to interpret.^7,21^ Although each field of a MACCS fingerprint corresponds to meaningful chemical properties (such as whether the chemical contains multiple aromatic rings, or at least one nitrogen atom), the fingerprint is largely inscrutable in QSAR applications, since biological activity is the result of many higher-order interactions between the chemical of interest and biomolecules.

In this study, the knowledge representation-based heterogeneous graph data represent easily interpretable relationships between entity types that mediate toxic responses to chemicals. Although not implemented in this particular study, a GNN architecture known as a *graph attention network* explicitly highlights portions of a graph that are influential in predictions, providing a logical next step for continuing this body of work on GNNs in QSAR modeling. Other, simpler approaches also provide avenues for exploring interpretability, such as visualizing the edge weights for edges starting at assay nodes in the trained GNN. Often, the sheer size of graphs make this approach intractable, but since our graph only contains 52 assays it is relatively straightforward to inspect their weights. For example, the highest weighted assays for the HepG2 cell viability prediction task are *HepG2 Caspase-3/7 mediated cytotoxicity* and *NIH/3T3 Sonic hedgehog antagonism* (a marker of developmental toxicity). The first of these makes sense from an intuitive standpoint, as it measures toxic response in the same cell line as the predicted assay. The second, on the other hand, does not have an immediately obvious connection to the predicted assay, but may be linked to the fact that Shh antagonists can induce apoptosis.^22^ Either way, it is easy to see that assay weights can be used to generate specific hypotheses for future targeted studies of mechanisms that underlie toxicity.

We provide all assay weights for the two above-mentioned assays in **Supplemental Materials**.

### 4.3. Sources of bias and their effects on QSAR for toxicity prediction

Like any meta-analysis technique, QSAR is subject to multiple sources of bias that can be introduced at several levels, not the least of which is in the original experiments used to generate toxic activity annotations for training data samples. This was a greater issue historically, when known activities for chemicals were compiled either from published scientific journal article results or from reporting guidelines for *in vivo* experiments.^23^ Publication bias caused negative activity annotations to be extremely incomplete, and techniques for imputing negative annotations were inconsistent. Older QSAR studies often did not state the original sources of their data, so verification and reproducibility of results are immensely challenging (if not impossible).

Fortunately, modern large-scale screening efforts (including Tox21) were created to directly address these and other issues.^24^ While our training data are still subject to batch effects, bias in selecting assays and chemicals for screening, and other systematic and experimental errors that are propagated along to the final QSAR model, we are relatively confident that publication bias, reporting bias, and other issues that plagued early QSAR studies have been substantially decreased. Furthermore, our GNN approach to QSAR modeling may be more robust to these sources of bias than non-GNN approaches, because (a.) the graph incorporates multiple levels of biological knowledge that can ‘fill in gaps’ left by incomplete or inaccurate data at other levels and (b.) GNNs—and heterogeneous GNNs in particular—exhibit properties that make them inherently robust to noise.^25,26^

## 5. Conclusions

In this study, we introduce a novel GNN-based approach to QSAR modeling for toxicity prediction, and evaluate it on data from 52 assays to show that it significantly outperforms existing methods. GNNs comprise an incredibly active emerging topic within artificial intelligence research, and as one of the first GNN applications in computational toxicology we hope that our results serve as a ‘jumping off point’ for a vast body of similar work. We plan to evaluate graph attention networks, new data modalities, and network regulization techniques in the near future, and encourage contributions from the toxicology and informatics communities at-large to improve predictive toxicology’s overall data ecosystem.

## 6. Code availability

All source code pertaining to this study is available on GitHub at https://github.com/EpistasisLab/qsar-gnn. A frozen copy of the code at the time of writing is available at https://doi.org/10.5281/zenodo.5154055.

## 7. Supplemental Materials

Supplemental tables and figures are available on FigShare at https://doi.org/10.6084/m9.figshare.15094083.

## Acknowledgements

This work was made possible with support from US National Institutes of Health grants `R01-LM010098, R01-LM012601, R01-AI116794, UL1-TR001878, UC4-DK112217` (PI: Jason Moore), `T32-ES019851`, and `P30-ES013508` (PI: Trevor Penning).

## Appendix A. Graph convolutional network architecture

Our GCNN implementation uses a message-passing paradigm that combines aspects of the GraphSAGE^27^ and R-GCN^16^ architectures. Let 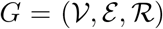 be a heterogeneous graph consisting of nodes 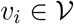, edges 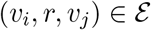 and a set of *edge types* 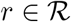. Each edge is labeled with exactly one edge type. All chemical nodes (represented below as *h*^0^) are represented by a bit string of length 166 corresponding to the chemical’s MACCS fingerprint, while all other nodes (assays and genes) are represented by a single decimal-valued ‘embedding feature’ that is learned during optimization. The magnitude of an assay or gene’s embedding is roughly equivalent that node’s importance in the network, and can be introspected for model interpretation.

Each layer of the network is defined as an edge-wise aggregation of adjacent nodes:

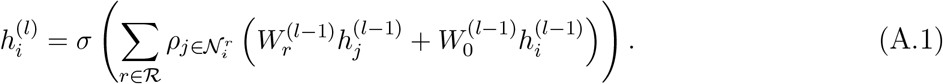

where 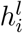 is the hidden representation of node *i* in layer *l*, 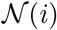 is the set of immediate neighbors of node *i*, and *σ* is a nonlinear activation function (either softmax or leaky ReLU, as explained in **Appendix B**). *ρ* can be any differential ‘reducer’ function that combines messages passed from incident edges of a single type; in the case of this study we use summation. Since our graph contains relatively few edge types, regularization of the weight matrices *W* is not needed.

## Appendix B. Node classification model

For classifying chemicals as active or inactive with regards to an assay of interest, we stack 2 GCN layers in the form given by (A.1), with a leaky ReLU activation between the two layers and softmax applied to the second layer’s output. Since we only classify chemical nodes, we ignore outputs for all other node types (and for chemicals with undefined labels); labels are generated via **Algorithm 1** We train the network by minimizing binary cross-entropy between the network’s softmax outputs and true activity values:

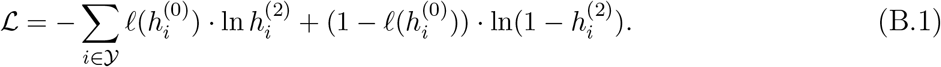

where 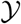 is the set of all labeled nodes, 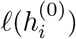 layer output of node *i*. is the true label of node *i*, and 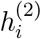 is the final layer output of node *i*.

The relatively shallow architecture of the network allows us to optimize the model using the Adam algorithm applied to the entire training data set, but the model can be adapted to mini-batch training when appropriate or necessary.

## Footnotes

a The full graph database for ComptoxAI can be found at https://comptox.ai, and will be described in a separate, upcoming publication.

b To prevent information leakage, since conectivity to the assay would perfectly predict the node labels.

